# Molecular Evolution of SARS-CoV-2 Structural Genes: Evidence of Positive Selection in Spike Glycoprotein

**DOI:** 10.1101/2020.06.25.170688

**Authors:** Xiao-Yong Zhan, Ying Zhang, Xuefu Zhou, Ke Huang, Yichao Qian, Yang Leng, Leping Yan, Bihui Huang, Yulong He

**Author notes:** Correspondence authors: Bihui Huang, Address: No.628, Zhenyuan Road, Guangming District, Shenzhen 518107, China, Yulong He, Address: No.628, Zhenyuan Road, Guangming District, Shenzhen 518107, China.

## Abstract

SARS-CoV-2 caused a global pandemic in early 2020 and has resulted in more than 8,000,000 infections as well as 430,000 deaths in the world so far. Four structural proteins, envelope (E), membrane (M), nucleocapsid (N) and spike (S) glycoprotein, play a key role in controlling the entry into human cells and virion assembly of SARS-CoV-2. However, how these genes evolve during its human to human transmission is largely unknown. In this study, we screened and analyzed roughly 3090 SARS-CoV-2 isolates from GenBank database. The distribution of the four gene alleles is determined:16 for E, 40 for M, 131 for N and 173 for S genes. Phylogenetic analysis shows that global SARS-CoV-2 isolates can be clustered into three to four major clades based on the protein sequences of these genes. Intragenic recombination event isn’t detected among different alleles. However, purifying selection has conducted on the evolution of these genes. By analyzing full genomic sequences of these alleles using codon-substitution models (M8, M3 and M2a) and likelihood ratio tests (LRTs) of codeML package, it reveals that codon 614 of S glycoprotein has subjected to strong positive selection pressure and a persistent D614G mutation is identified. The definitive positive selection of D614G mutation is further confirmed by internal fixed effects likelihood (IFEL) and Evolutionary Fingerprinting methods implemented in Hyphy package. In addition, another potential positive selection site at codon 5 in the signal sequence of the S protein is also identified. The allele containing D614G mutation has undergone significant expansion during SARS-CoV-2 global pandemic, implying a better adaptability of isolates with the mutation. However, L5F allele expansion is relatively restricted. The D614G mutation is located at the subdomain 2 (SD2) of C-terminal portion (CTP) of the S1 subunit. Protein structural modeling shows that the D614G mutation may cause the disruption of salt bridge among S protein monomers increase their flexibility, and in turn promote receptor binding domain (RBD) opening, virus attachment and entry into host cells. Located at the signal sequence of S protein as it is, L5F mutation may facilitate the protein folding, assembly, and secretion of the virus. This is the first evidence of positive Darwinian selection in the *spike* gene of SARS-CoV-2, which contributes to a better understanding of the adaptive mechanism of this virus and help to provide insights for developing novel therapeutic approaches as well as effective vaccines by targeting on mutation sites.

## Introduction

Severe acute respiratory syndrome coronavirus 2 (SARS-CoV-2), the causative agent of an emerging coronavirus disease (COVID-19) that has caused more than430,000 deaths, is still a serious global pandemic currently. The genome of SARS-CoV-2 is consisting of a single-stranded and positive-sense RNA of around 30 kb in length with a 5’ cap and 3’-polyA tail. It shows that SARS-CoV-2 genome possesses six major open reading frames (ORFs) that encodes 27 different proteins, in which four are structural proteins named Envelope (E), Membrane (M), Nucleocapsid (N) and Spike (S). Many studies have demonstrated important functions of these proteins in virus entry, transcription and virion particle assembly of SARS-CoV-2. The E protein is a small envelope protein with 75 amino acids. Given that a close genetic relationship between SARS-CoV-2 and SARS-CoV, functions of this protein may include virion assembly and morphogenesis[1]. In addition, induction of apoptosis of host cells might be another crucial function of SARS-CoV-2 E protein, thus making it a potential determinant of viral pathogenesis [2]. M protein, consisting of 222 amino acids, is the most abundant component of the viral envelope and plays a key role in the virion assembly[3]. N protein, composed of 419 amino acids, may form complexes with genomic RNA, interact with the viral membrane protein, and play a critical role in enhancing the efficiency of virus transcription and assembly[4]. S protein, consisting of 1,273 amino acids, is the most important factor that mediates virus entry and a primary determinant of cell tropism and pathogenesis of SARS-CoV-2[5].

Many studies demonstrated SARS-CoV-2 underwent the evolution and some genetic evolutionary features have been reported[6]. The whole genomic sequence of SARS-CoV-2 has 79.6% identity with SARS-CoV and 96% with a bat SARS-related coronavirus (SARSr-CoV), RaTG13. Although no positive time evolution signal was found between SARS-CoV-2 and RaTG13, the SARS-CoV-2 shows a strong positive temporal evolution relationship with bat-SL-CoVZC45, which has a slightly less identical genomic sequence (87.5%) than RaTG13 [7]. Combining the phylogenetic analysis of full-length genomes of coronaviruses, a potential bat origin of SARS-CoV2 is indicated [8]. A recent study reported that *spike* (S) gene (coding gene of S protein) of SARSr-CoVs from their natural reservoir host, the Chinese horseshoe bat (*Rhinolophus sinicus*), has coevolved with *R. sinicus* angiotensin converting enzyme 2 (ACE2) via positive selection[9]. A single-stranded positive-sense RNA virus as it is, SARS-CoV-2 causes global pandemic within half a year, suggesting it may evolve rapidly. However, the evolution of SARS-CoV-2 based on structural genes from human to human transmission has not been investigated in detail. The primary purpose of this work is to study the evolutionary pattern of the four structural genes of SARS-CoV-2 derived from a global isolate collection including the E, M, N and S. Various molecular evolution and selection analysis approaches were employed to identify the phylogeny of the four structural proteins and potential selection effects on these genes. Hereby, our study reveals that intragenic recombination does not contribute to the evolution of these genes while purifying selection is the main evolutionary force. Moreover, a D614G mutation in the S protein is operated by strong positive selection and may be responsible for the quick spread of SARS-CoV-2 globally. Additionally, another potential L5F mutation may also be operated by positive selection, but with relatively less strong pressure as compared to D614G.

## Materials and Methods

### SARS-CoV-2 isolates

Complete full-length genomic sequences of SARS-CoV-2 were downloaded from 2019 Novel Coronavirus Resource (2019nCoVR) in China National Center for Bioinformation. All of which were also uploaded to the NCBI GenBank database. The sequences were manually checked and finally a total of 3090 isolates were selected and verified for the present study. These isolates were collected from December 24, 2019 to April 24, 2020 in the different geographical locations including China, USA, Japan, Pakistan, Australia, Greece, German, Peru, Turkey, Kazakhstan, Iran, Serbia, Thailand, Nederland, Sri Lanka, Czech, Malaysia, India etc. Detailed information of these isolates including the GenBank accession number or biosample number is summarized in S1 Table.

### Sequence analysis of the four structural genes and proteins

The E, M, N, S gene sequences were extracted from SARS-CoV-2 global isolate collection and aligned by the MEGA X package using Muscle (codons) parameters [10]. Because some regions of genomic sequences of SARS-CoV-2 couldn’t be exactly identified, in which nucleic acid bases are shown as degenerate bases (e.g. N, R, Y), we were unable to obtain all of the four structural gene sequences from an isolate sometimes. Allele type and DNA sequence polymorphism analyses were performed using DnaSP 6.12.03[11]. The protein sequences and polymorphism loci of these isolates were also aligned and analyzed with the MEGA X.

### Molecular evolution analysis

An unrooted phylogenetic tree of the four structural proteins was constructed using the MEGA X package [10], and the evolutionary history was inferred using the Maximum Likelihood method, based on the JTT matrix-based model for E protein sequences, General Reversible Chloroplast + Freq. model for M, JTT matrix-based model for N and Jones et al. w/freq. model for S protein sequences. Model selection was conducted in MEGA X. Bootstrap values were estimated by 1000 replications. Initial tree(s) for the heuristic search were obtained automatically by applying Neighbor-Join and BioNJ algorithms to a matrix of pairwise distances estimated using each model mentioned above. The tree is drawn to scale, and FigTree V1.4 was utilized to form cladogram branches (http://tree.bio.ed.ac.uk/software/figtree/). The aligned DNA sequences were also screened using RDP4 software to detect intragenic recombination among the alleles of each structural gene[12]. Six methods implemented in the RDP4 were utilized. These methods are RDP [12], GENECONV[13], BootScan [14], MaxChi[15], Chimaera [16], and SiScan [17]. Common settings for all methods include considering sequences as linear and setting statistical significance at the P < 0.05 with Bonferroni correction for multiple comparisons and requiring phylogenetic evidence and polishing of breakpoints. Potential recombination events (PREs) were considered as those identified by at least two methods. Reticulate network tree of alleles of the four structural genes of SARS-CoV-2 was also generated by Splitstree4 [18]. Phi test implemented in Splitstree4 was used to define probable recombination events. Tajima’s D, Fu and Li’s D* and F* tests were employed to test the mutation neutrality hypothesis of the whole gene as previously described by our research group[19]. These analyses were carried out using DnaSP 6.12.03[11]. A statistical significance level with P < 0.05 is acceptable. The false discovery rate and 1000 replications in a coalescent simulation were applied for correcting multiple comparisons. Non-neutrality evolution was considered when identified by at least two out of three tests. Nonsynonymous and synonymous mutations of the alleles of the four structural genes were also calculated using MEGA X package [10].

### Analysis of positive selection based on codon

The selection pressure operating the four structural genes of SARS-CoV-2 was searched by using the Maximum Likelihood (ML) method. Analyses were performed using a visual tool of codeml program, named EasyCodeML algorithm with site model [20]. Three nested models (M3 vs. M0, M2a vs. M1a, and M8 vs. M7) were compared and likelihood ratio tests (LRTs) were applied to access a better fit of codes. Model fitting was also performed using multiple seed values for *dN/dS* and assuming the F3×4 model of codon frequencies. Positive selection is inferred when individual site or codon with ratio of nonsynonymous to synonymous mutations (*dN/dS* ratios) is greater than one (ω>1). When the LRT is significant (*p <0.05*), Bayes empirical Bayes (BEB) (M8 model) and Naive Empirical Bayes (NEB) methods (M3 and M2a model) are further employed to identify amino acid residues that likely evolve under positive selection based on a posterior probability threshold of 0.95. Results from M8 model were taken as the standard as Yang *et al.* reported. M3 model was used for the frequency distribution of codon class analysis as Yang *et al.* recommended[21]. HyPhy package was used to validate the result obtained by ML method[22].

### Structural modeling of the protein with positive selection sites

Three-dimensional structures of proteins with positive selection sites were modeled using SWISS-MODEL (http://swissmodel.expasy.org) according to the most fitted protein template. Model quality was evaluated by QMEAN while the structure of the model was visualized by using PyMoL [23].

## Results and Discussion

### Characteristics of SARS-CoV-2 isolates, structural gene and protein sequences

The 3090 SARS-CoV-2 isolates harbor only 16 unique alleles of E and 40 alleles of M, but an abundant number of alleles of N and S genes, which contain 131 and 173, respectively. These alleles correspond to 10, 14, 88 and 99 different amino acid sequences of E, M, N, and S proteins, respectively. Protein sequence comparisons of WH01 isolate with SARSr-CoV, bat-SL-CoVZC45 isolate show 100% (75/75) identity in E, 98.65% (219/222) identity in M, 94.27% (395/419) identity in N and 80.06% (1171/1273) in S proteins, respectively. These results imply a close kinship between SARS-CoV-2 and bat SARSr-CoV, especially on E and M proteins. On the other hand, it indicates an extreme conservation of E and M proteins and their functions among coronaviruses[24].

Further analysis revealed that there are 14 single nucleotide polymorphisms (SNPs) of E gene, but only 5 single amino acid polymorphic (SAP) loci in the E protein. Similar result was observed on M gene and protein, with 37 SNPs and 9 SAPs. In contrast, 126 SNPs and 75 SAPs are detected on N gene and protein, respectively. S protein, the most important factor that mediates virus entry by receptor binding and membrane fusion and determines the infection ability of SARS-CoV-2 [25], harbors 155 SNPs on the alleles and 90 SAPs in the protein. Considering the size of nucleotides and amino acid residues, N gene has the maximum sequence variability with 10.02% (126/1257) SNPs and 17.90% (75/419) SAPs, respectively. However, S gene has most pairwise nucleotide differences among the four structural genes, indicating a more genetic diversity of S gene (Table 1). A key player in the virus transcription and assembly as N protein is [26, 27], high sequence variability of the N protein may indicate a vast adaption of the virus during host transmission. Previous study shows that high genetic variance has been found among bat SARSr-CoVs, particularly in the S gene[9]. Similar, higher nucleotide diversity (π, a major parameter to define genetic diversity) of S gene is also detected on SARS-CoV-2 isolates, suggesting this may benefit virus survival in the host of human beings.

**Table 1.** Summary of genetic diversity of the 4 structural genes of the SARS-CoV-2 isolates.

| Gene | Sequence, n* | Sequence length | $h$ | $\pi$ | $S$ | $\theta$ | □ |
| --- | --- | --- | --- | --- | --- | --- | --- |
| E | 2928 | 228 | 16 | 0.00012 | 14 | 0.00475 | 15 |
| M | 2891 | 669 | 40 | 0.00018 | 37 | 0.00665 | 40 |
| N | 2253 | 1260 | 131 | 0.00056 | 126 | 0.01081 | 130 |
| S | 2339 | 3825 | 173 | 0.00075 | 155 | 0.00753 | 169 |

$h$ , Haplotypes,
$\pi$ , Nucleotide diversity
$S$ , Polymorphic sites
$\theta$ , Theta (per site) from $S$ , population mutation ration
□ , Total number of mutations
\* Some bases of SARS-CoV-2 genomic sequences are not exactly identified; thus, the number of gene sequences were less than 3090.

### Distinct phylogenetic patterns of the four structural genes

The phylogenetic analysis revealed that all SARS-CoV-2 E proteins form three clusters. Similar to E protein, phylogenetic tree of SARS-CoV-2 M proteins is formed by three clusters with few branches (Figs 1A and 1B). The results suggest both E and M genes may display a relatively high conservation during coronavirus evolution. In contrast, SARS-CoV-2 N and S proteins show distinct phylogenetic pattern as compared with that of E and M. Four and three main phylogenetic clusters with various branches are identified in the N and S proteins, respectively (Figs 1C and D). Given the crucial roles of N and S proteins in virus transcription, assembly, and entry to host cells, whether SARS-CoV-2 isolates harbor different N and S variants (such as those clustered into different clades) may influence their infection efficiency remains unknown, and requires further study.

**Figure 1.**
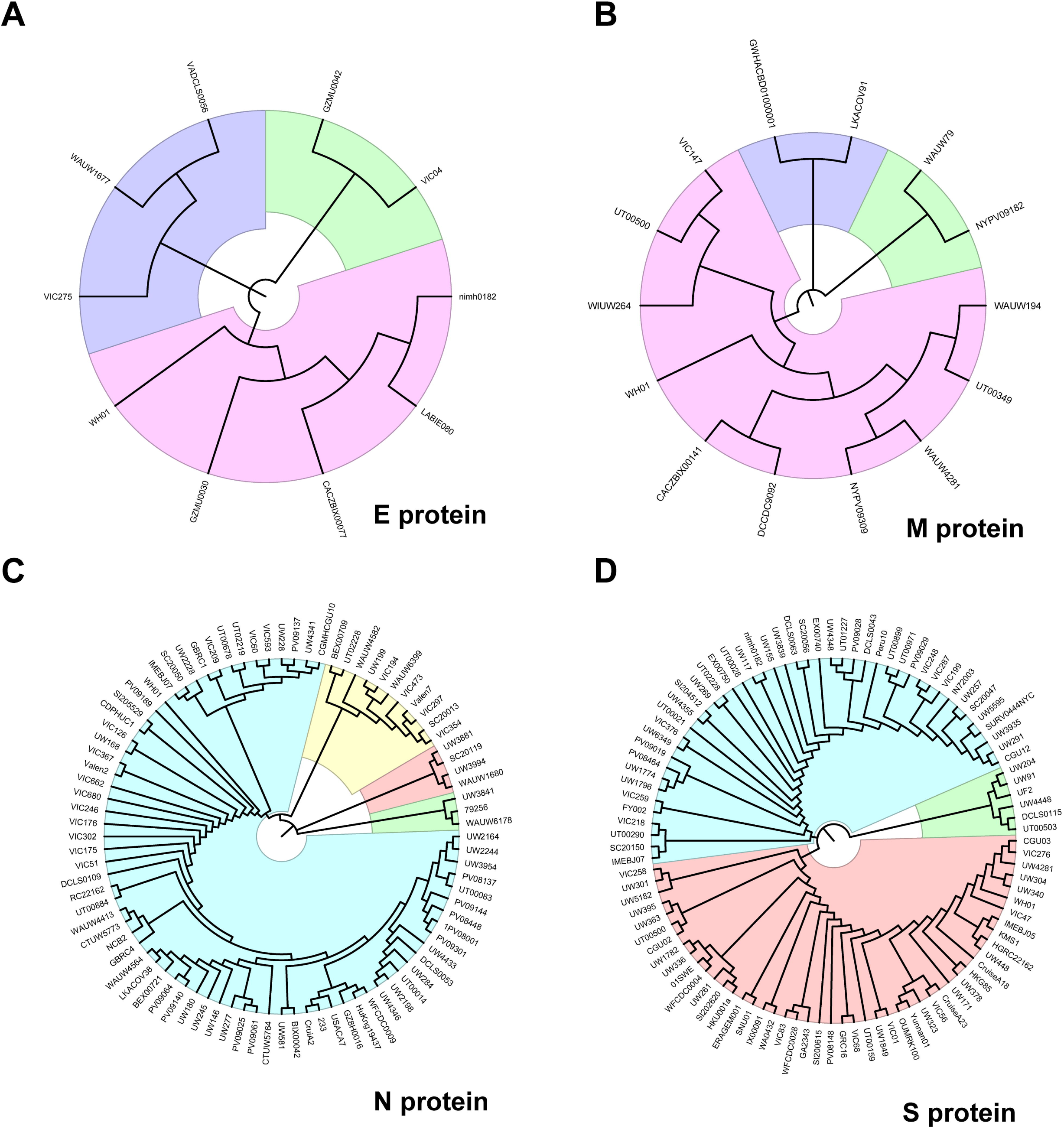
Phylogenetic tree of E (A), M (B), N (C), and S(D) proteins of SARS-CoV-2. Major clades are highlighted with different color. The tree shows topology of the protein of each allele, named by their representative isolates.

### Purifying selection drives the evolution at whole structural gene levels of SARS-CoV-2 during its human to human transmission

Although many studies demonstrated that recombination plays an important role on the emergence of SARS-CoV-2 and its contribution to admit SARS-CoV-2 as a human infectious pathogen [28-30], how this virus evolves during its global transmission has not been profiled yet. Therefore, we first analyzed intragenic recombination events of each structural gene using RDP4. The results indicate there were no recombination events occurred among the alleles of each gene (data not shown). Recombination event were also assessed through reticulate network tree by phi test in SplitsTree4. Although some internal nodes are noticed in N and S alleles, no significant evidence for recombination is validated of each gene by Phi test (p>0.05) (Fig 2). It indicates a relative stable state of SARS-CoV-2 during its transmission although a possible genetic interaction of different isolates might have occurred when it became a global pandemic [31, 32]. In addition, Tajima’s D, Fu and Li’s D* and F* statistics were calculated to examine the mutation neutrality hypothesis of the four structural genes of SARS-CoV-2. The results reveal that the evolution of all four genes does not match the neutral hypothesis, but favor purifying selection (Table 2 and Fig 3). The average of all pairwise *dN/dS* ratios (ω) among the alleles of each structural gene of SARS-CoV-2 is 0.5443 in E, 0.1562 in M, 0.07978 in N, and 0.4980 in S gene, respectively. All together, these results suggest that at the whole gene level, inconsistent purifying selection is the main evolution force (Table 2). Li et al studied the origin of SARS-CoV-2 and showed evidence of strong purifying selection in the S and other genes among bat, pangolin and human coronaviruses, indicating similar strong evolutionary constraints in different host species [33]. Similarly, our results suggest purifying selection drives the evolution at the whole structural gene level of SARS-CoV-2 during its transmission from human to human. This result also implies that in general, the genetic variation on these structural genes will not confer a significant disadvantage on the virus survival, and ratios reflect general variability of these genes and proteins. Considering that no recombination happened, nonsynonymous mutations would be removed at a great rate during the virus transmission [34].

**Table 2.** Summary of neutrality for the four structural genes in SARS-CoV-2 isolates.

| Gene | Tajima'sD | Fu and Li's D* test<br>t | Fu and Li's F<br>* test | dN | dS | dN/dS<br>($\omega$) | Selection |
| --- | --- | --- | --- | --- | --- | --- | --- |
| E | -2.29974, P<0.01 | -3.18477, P<0.02 | -3.38505, P<0.02 | 0.006836 | 0.1256 | 0.5443 | Purifying<br>selection |
| M | -2.74611, P<0.001 | -5.64276, P<0.02 | -5.50855, P<0.02 | 0.001294 | 0.008296 | 0.1562 | Purifying<br>selection |
| N | -2.87598, P<0.001 | -9.67153, P<0.02 | -7.95879, P<0.02 | 0.000251 | 0.003146 | 0.07978 | Purifying<br>selection |
| S | -2.87646, P<0.001 | -11.01171, P<0.02 | -8.59037, P<0.02 | 0.000609 | 0.001223 | 0.4980 | Purifying<br>selection |

**Figure 2.**
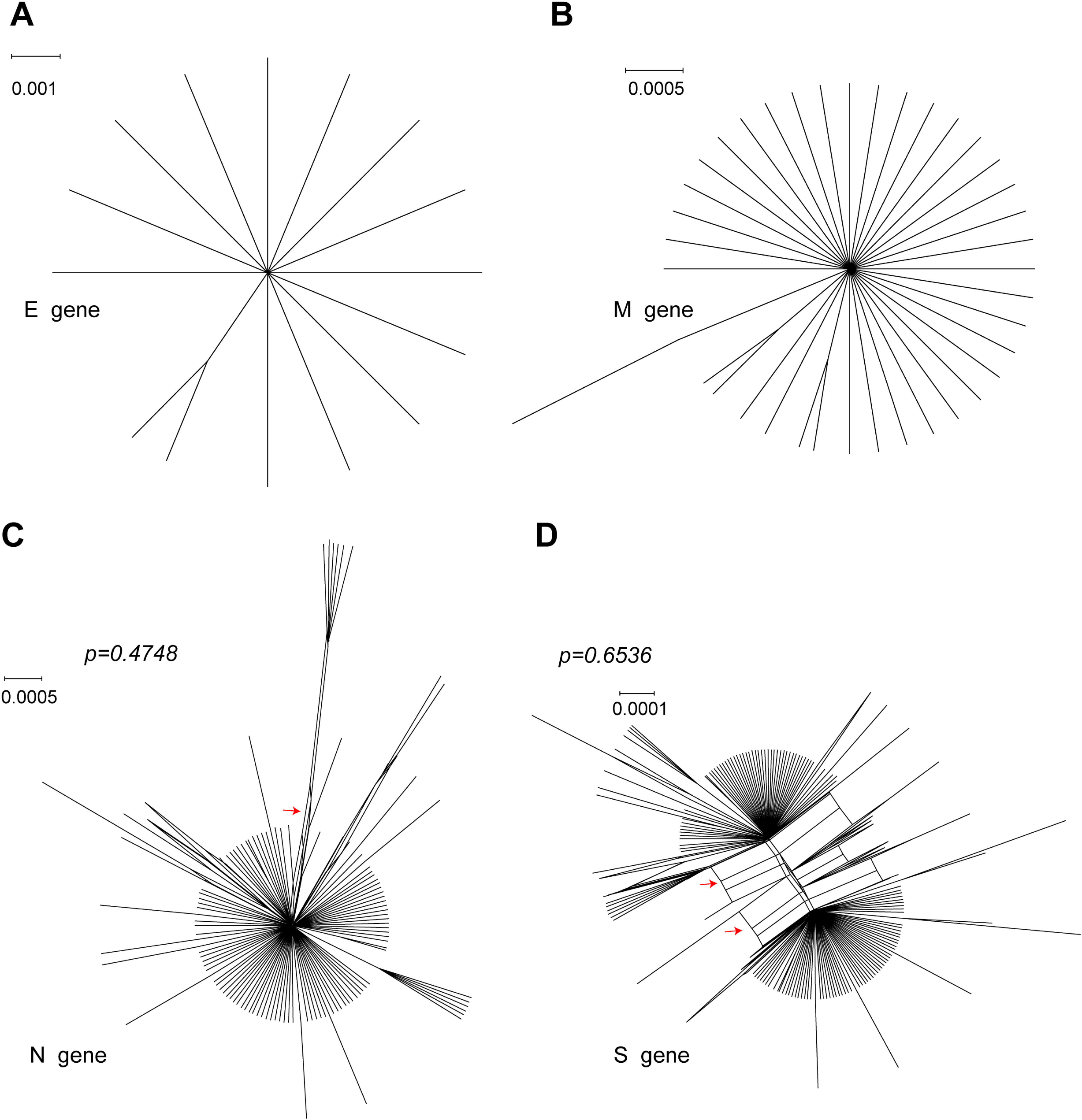
Reticulate network trees of E (A), M (B), N (C) and S (D) alleles of SARS-CoV-2 analyzed by the neighbor-net algorithm of SplitsTree4. Scale bars indicate number of substitutions per site. All internal nodes represent hypothetical ancestral alleles and edges that correspond to reticulate events such as recombination. Red arrows indicate edges. Because there are too few informative characters to use the Phi test for E and M genes, *p-values* of Phi test of N and S genes are shown.

**Figure 3.**
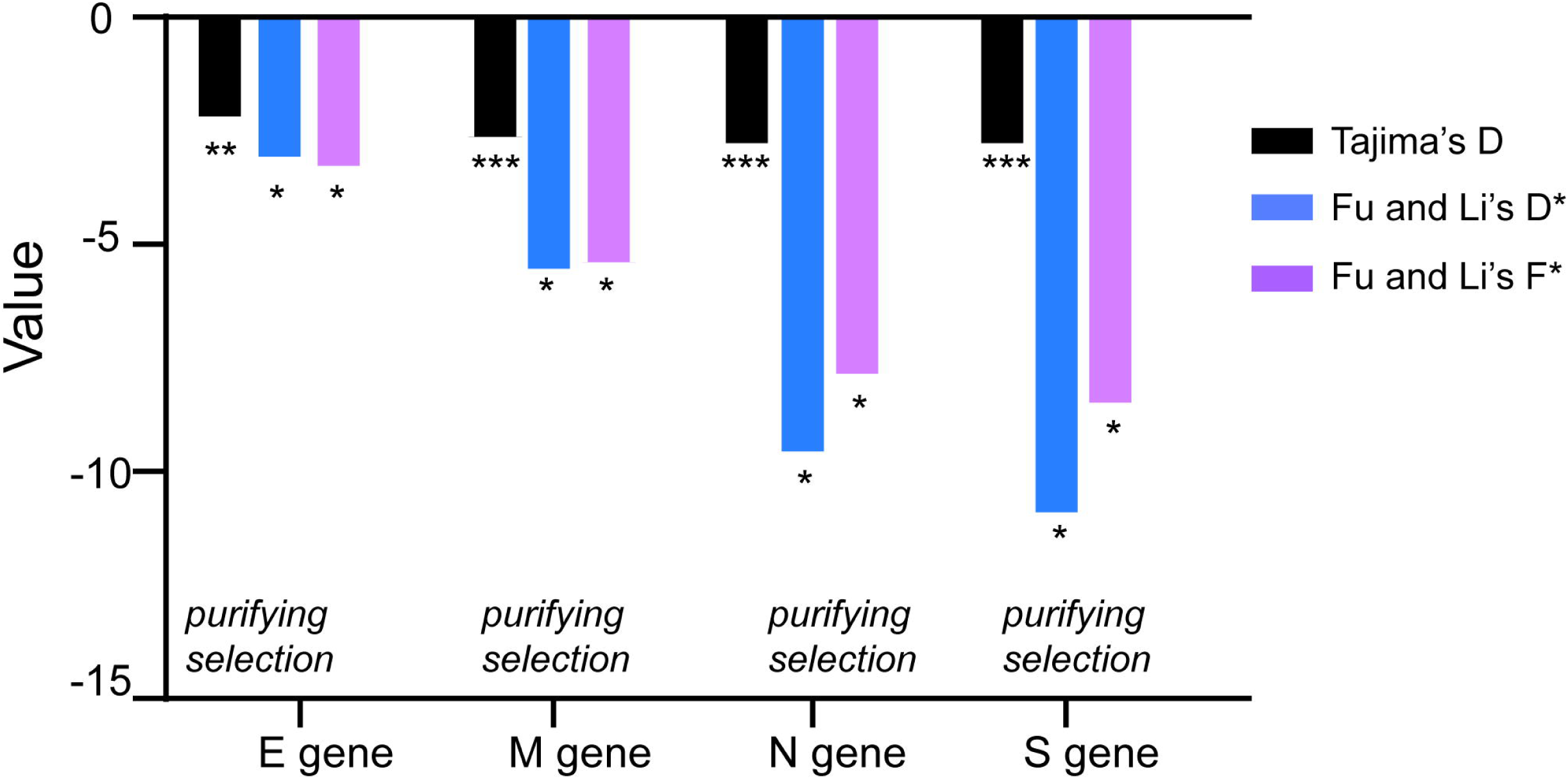
Tajima’s D, Fu and Li’s D* and F* test for the four structural gene alleles of SARS-CoV-2. *p < 0.05; **p < 0.01; ***p<0.001

### SARS-CoV2 S gene is operated by positive selection at a definitive codon located at the C-terminal portion of S1 subunit and a potential codon located at the signal sequence

Guo et al. reported that the S gene of SARSr-CoV populations in their natural host, Chinese horseshoe bat (*Rhinolophus sinicus*), has evolved through positive selection at some codons[9]. As mentioned above, at the whole gene level, purifying selection is the main force driving the evolution of studied genes. Whether positive selection pressure accelerates the diversification of the structural genes of SARS-CoV-2 remains unclear. Therefore, we used codon-substitution models to estimate the ratio of nonsynonymous over synonymous substitutions (*dN/dS*), also known as ω. The role of recombination in the polymorphism of four genes is excluded because no intragenic recombination was detected (Fig 2). By using ML model, we don’t find any codon of E and M gene subjecting to positive selection obviously (data not shown). However, a potential positive selection site 208A in N gene is identified by using M3 model, but not by any other models especially the M8 model, suggesting a limited amount of evidence of positive selection in N gene (S1 Table). For the S gene, we found the average ω is 0.37199 calculated by M0 model of the codeML package, suggesting that purifying selection was a major force operating the evolution of the S gene during its transmission among human beings. In three LRTs, all alternative models (M3, M2a, M8) are significantly better fit (P<10^−4^) than relevant null models (M0, M1a, M7), indicating that some sites of S were subjected to strong positive selection (ω=18.22175-20.61283) (Table 3). A single positive selection site (614D) is identified in the S gene with posterior probability of 1.000 in all the three models [21], a clear evidence showing that this site is still experiencing positive selection when the virus transmitted from human to human. The result is also validated using internal fixed effects likelihood (IFEL) and Evolutionary Fingerprinting methods implemented in HyPhy package (Fig 4) [35-37]. To our surprise, the positive selection site is not located at the receptor binding domain (RBD) or receptor binding motif (RBM) as we anticipated, which play the most important role in virus-receptor interaction and virus entry into host cells [38]. This result suggests that a relatively genetic stability of this motif would benefit the virus survival. Intriguingly, the site under positive selection pressure always has a D614G (for the S gene is 1841A>G) mutation, implying such mutation may enhance virus adaptability in human hosts. Another potential positive selection site at codon 5 is also identified, and a L5F mutation (for the S gene is 13C>T) is always found, with posterior probabilities greater than 0.95, 0.93 and 0.92 (critical values) calculated by M3, M2a and M8 models (Table 3), respectively. Similar result was also confirmed by Evolutionary Fingerprinting method (S1 Fig). Considering signal sequence (SS) is a short hydrophobic peptide that plays an important role in guiding viral protein into the endoplasmic reticulum (ER) for proper folding and assembly [39], we postulate that L5F mutation may increase hydrophobicity of the SS, thus facilitating the entry of S protein into ER for folding and assembly, and in turn secretion of the virus.

**Table 3.** Log-likelihood values and parameter estimates for the SARS-CoV-2 S gene sequences.

| Model | Ln L | Estimates of parameters | Model compared | LRT P-value | Positive sites |
| --- | --- | --- | --- | --- | --- |
|  |  | p0=0.96797, p1=0.02883, p2=0.00320 |  |  | **5 L 0.958\***,28 Y 0.850,221 S 0.901,**614 D** |
| M3 (discrete) | -6766.339162 | ω0=0.26126, ω1= 2.70530, **ω2=20.61283** | | | **1.000\*\*\***,677 Q 0.891 |
| M0 (one ratio) | -6790.072925 | ω0=0.37199 | M0 vs. M3 | 0.000000001 | Not Allowed |
|  |  | p0=0.81731, p1=0.17872, p2=0.00397 |  |  | 5 L 0.9258,28 Y 0.812,221 S 0.832,**614 D** |
| M2a(selection) | -6766.432802 | ω0=0.17504, ω1=1.00000, **ω2=18.76936** | | | **1.000\*\*\***,677 Q 0.828 |
|  |  | p0=0.70461, p1=0.29539 |  |  |  |
| M1a (neutral) | -6778.770190 | ω0=0.04395, ω1=1.00000 | M1a vs. M2a | 0.000004385 | Not Allowed |
|  |  | p0=0.99578, p=0.40368, q=0.82224 |  |  | 5 L 0.931,28 Y 0.817,221 S 0.831,**614 D** |
| M8(beta&$\omega$) | -6768.829411 | p1= 0.00422, **$\omega$= 18.22175** | | | **1.000\*\*\***,677 Q 0.828 |
| M7(beta) | -6779.230494 | p=0.00857, q=0.02623 | M7 vs.M8 | 0.000030400 | Not Allowed |

LnL is the log likelihood; $\omega$ is ratio of $dN/dS$ , LRT P-value indicates the value of chi-square test; Parameters indicating positive selection are presented in bold;
Positive selection sites were identified by the Bayes empirical Bayes (BEB) methods under M8 model. The posterior probabilities (p) ≥ 0.80 are shown, (p) ≥ 0.95
(p) ≥ 0.99, and (p)=1.000 are indicated by \*, \*\* and \*\*\*, respectively. Yang *et al.* recommended that results from M8 model were preferred to find sites under
positive selection pressure.

**Figure 4.**
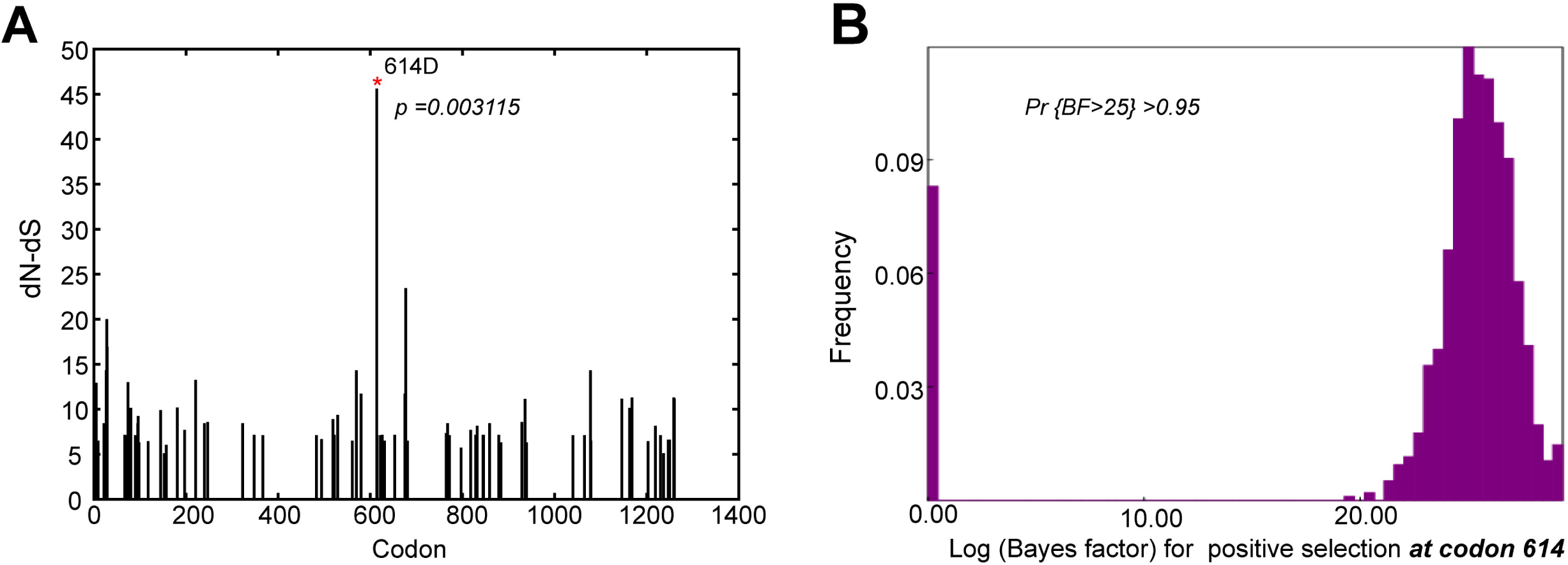
Positive selection analysis of S gene codons by IFEL and Evolutionary Fingerprinting methods. **A.** Diagram of selection analysis result of S codons by IFEL method. Asterisk indicates the positive selection site with statistical significance (*p<0.01*). **B.** Log (Bayes Factor) for positive selection at codon 614 of S gene and its frequencies. The cut-off value for the Bayes factor (BF) in the Evolutionary Fingerprinting method was set at 25 to reflect a positive selection at a given site (Posterior probability>0.95). Pr {BF>25} indicates posterior probability of Bayes Factor >25.

### Evolutionary relationship of S gene alleles with or without D614G and L5F mutation

Phylogenetic tree of S gene alleles was derived to test the evolutionary relationship among the alleles with or without D614G mutation. As shown in Fig 5A, the 173 alleles of the S gene could be clustered into four clades. Alleles with D614G mutation could be found in all 4 clades, among which a dominant one contains 79 out of 85 alleles with such mutation. The remaining 6 mutated S alleles are distributed in other 3 clades. The result suggests a potential common ancestor for the majority of S alleles with D614G mutation, while some other maybe derived from alternative ancestors. This result is also supported by the parsimony network of S gene alleles using PopART (http://popart.otago.ac.nz) [40]. Two central alleles (representative virus isolates are WH01 and GZMU0019) and associated alleles around them form a star scattering network, suggesting that the S gene may have two potential origins (Fig 5B). All S alleles with D614G mutation are closely related (with a few point mutations), and comprise a scattered star structure, suggesting the expansion of SARS-CoV-2 population with D614G mutation on S gene. In contrast, alleles of the N gene show a single ancestor analyzed by parsimony network though 3 phylogenetic clades are identified (S2 Fig).

**Figure 5.**
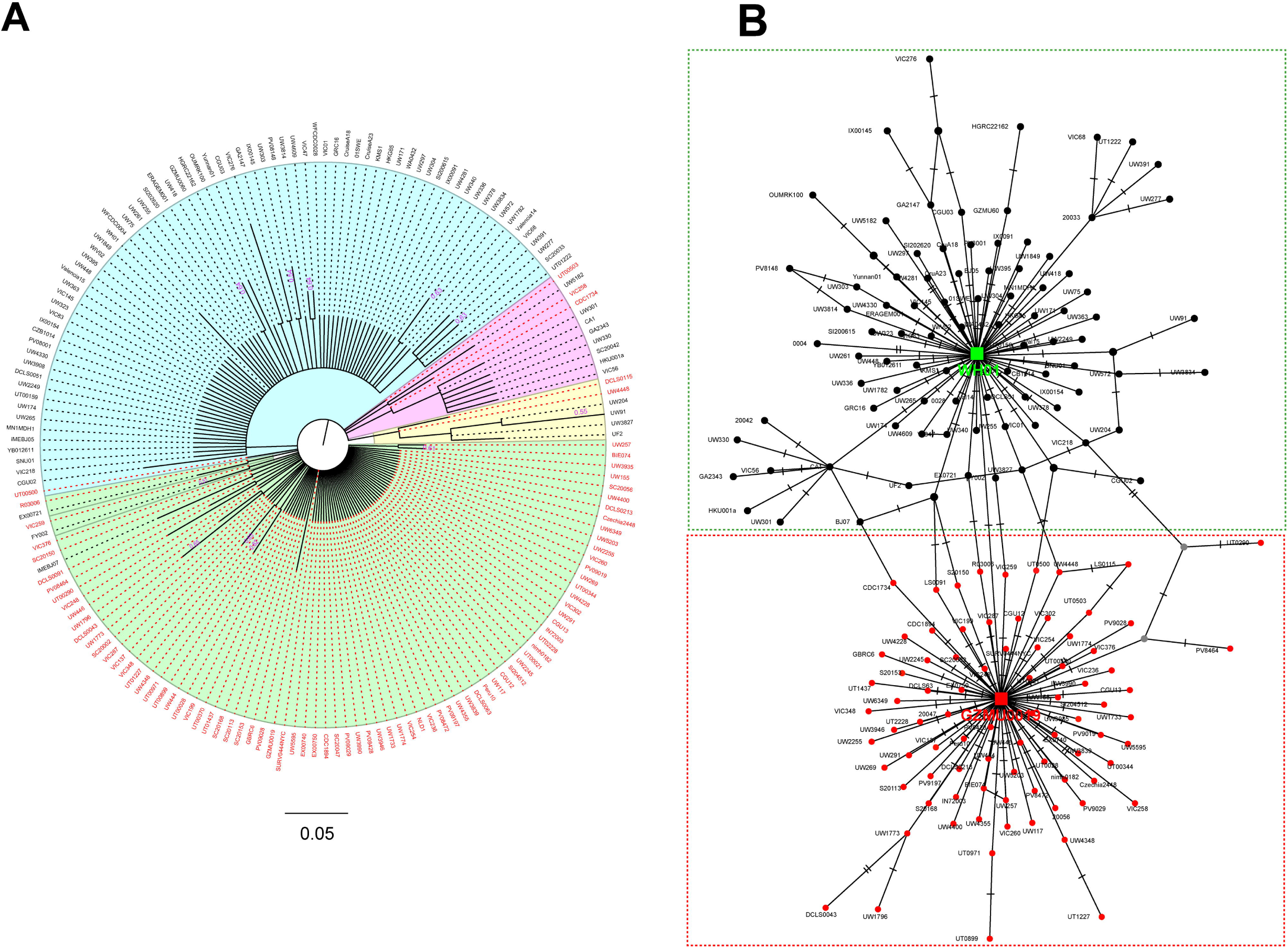
Evolutionary relationship of S alleles with or without D614G mutation. **A.** Phylogenetic tree of S gene based on nucleotide sequences of 173 alleles. The evolutionary history is inferred using the Maximum Likelihood method and Tamura-Nei model. The tree is drawn to scale, with branch lengths measured in the number of substitutions per site. Each clade is highlighted with different color. Alleles are shown with their representative isolate names, and alleles with D614G mutation are highlighted in red. Bootstrap values more than 0.5 are shown. **B.** Parsimony network of SARS-CoV-2 S gene haplotype (allele) diversity obtained from 3090 isolates worldwide. Each oblique line linking between haplotypes (haplotype name is shown as its representative isolate name) represents one mutational difference. Unlabeled nodes (Gray circle) indicate inferred steps have not found in the sampled populations yet. The ancestral haplotype, or root of the network, is labeled with a square, and represent haplotype name is marked green or red. The red nodes indicate haplotypes with D614G mutation, while green or black nodes indicate haplotypes without D614G mutation. Dotted boxes indicate major haplotype groups.

A total of 5 alleles with L5F mutation are found and all of them are in one clade, accounting for 83.33% of all alleles in the clade (S3A Fig). Further parsimony network analysis reveals that S alleles with L5F mutation are not closely related, but distribute in both WH01 and GZMU0019 haplotype groups (S3B Fig). No scattered star structure of these alleles can be formed, indicating L5F mutation might arise from independent origins other than that of D614G mutants. Limited number of alleles with L5F mutation identified so far also suggests that L5F might subject to relatively less strength of the pressure and is still at early stage of positive selection.

### Frequency of S allele with D614G mutation increased in SARS-CoV-2 isolates during human to human transmission

Considering that mutation of a positive selection site should be beneficial to the survival of the individuals carrying the mutation, we postulate that the D614G (1841A>G) mutation may help the spread of SARS-CoV-2. Some evidence has been obtained from the haplotype network of S alleles mentioned above (Fig 5B). S gene haplotypes (alleles) with D614G mutation (representative isolate GZMU0019) have evolved many subtypes and comprise a star structure with GZMU0019 in the center. This starburst pattern with one haplotype in the center and many other haplotypes surrounding the central haplotype suggests a signature of rapid population expansion [41]. To further study whether SARS-CoV-2 isolates with D614G mutation have advantage in survival during its transmission among human beings, we calculated the frequencies of S alleles carrying D614G mutation in each week from the collected SARS-CoV-2 isolates from December 24, 2019 to April 20, 2020 (17 weeks). Detailed information of these isolates including collection date, collection region and accession or biosample numbers is summarized on S3 and S4 Tables.

In 173 S gene alleles, 85 carry D614G mutation, accounting for 49.13% of all. Similarly, 47 out 99 S proteins carry D614G mutation, accounting for 47.47% of all. The first two isolates, GWHABKF00000001 and WH01 (isolated in December 24, 2019 and December 26, 2019, respectively), carry 614D in the S protein, while the first SARS-CoV-2 isolate with a D614G mutation is GZMU0019 in our collected dataset, isolated from a patient with COVID-19 on February 5, 2020 (week 7 in our dataset). After that, except for week 9 and week 10 (possibly due to the small number of samples and sampling deviation), a spread trend that more and more proportion of isolates carry the D614G mutation in the S protein stands out. In the week 17, the last week of our dataset, 91.11% of SARS-CoV-2 isolates carry this mutation (S3 Table, Fig 6A). Further analysis reveals that the frequency of D614G mutation in the S gene was steadily increasing when combining data from week 6 to 17 (S3 Table, Fig 6B). To exclude the influence of sample size on the result (in some weeks, only 4-6 isolates were collected in the dataset), we reorganized the dataset by taking both the sample size and sampling time into account. Various panels of 200-300 isolates were studied and similar results were observed (S4 Table, Figs 6C and D). Taken together, these results suggest that SARS-CoV-2 isolates with D614G mutation may increase their ability to transmit, and contribute to the rapid spread of this virus to the world.

**Figure 6.**
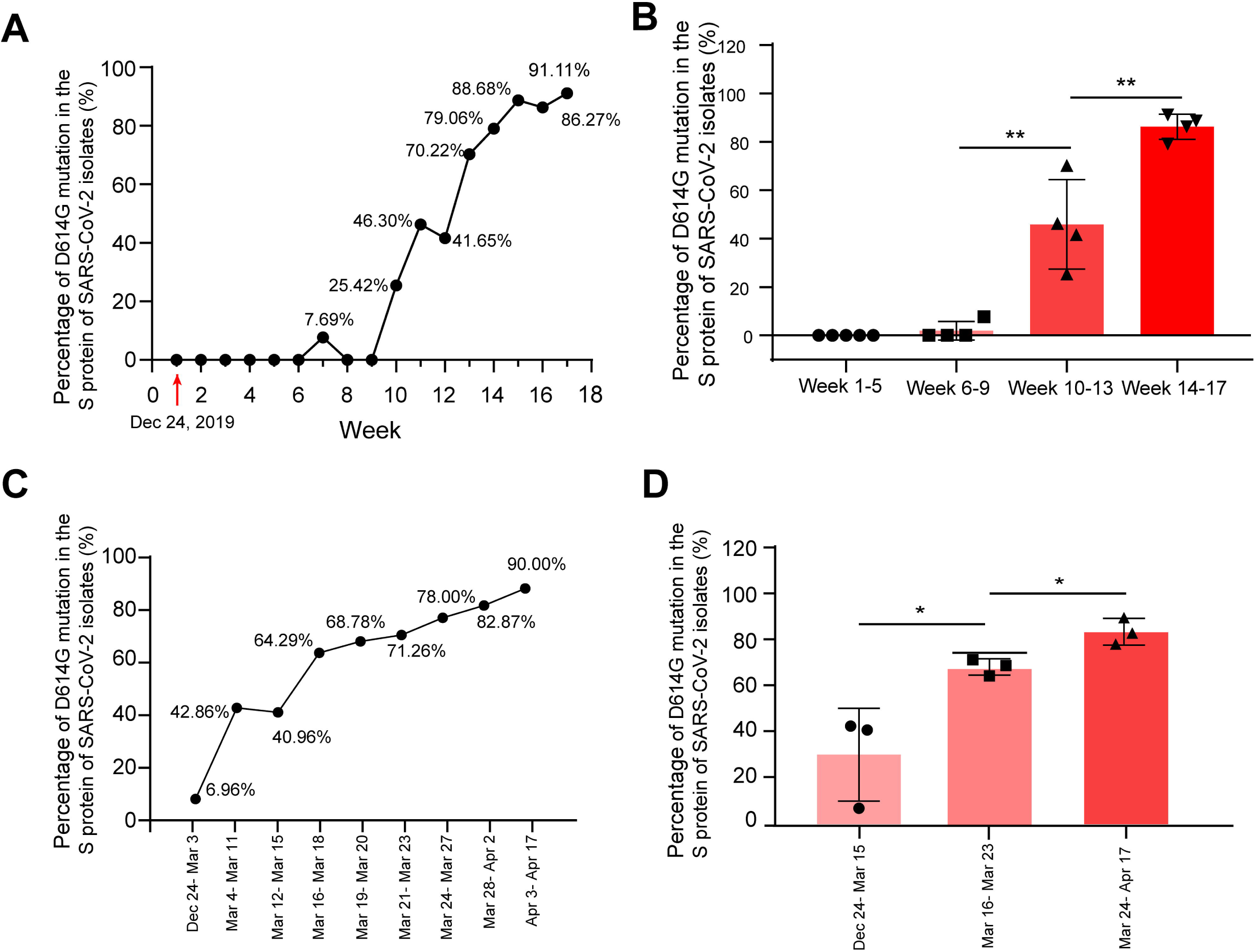
Expansion of S alleles with D614G mutation during SARS-CoV-2 human to human transmission. **A**. Percentage of SARS-CoV-2 isolates carrying the alleles of D614G mutation in each week collected. **B.** Frequencies of D614G mutation in the S gene in each period of time (Four to five weeks’ data are combined). **C.** Percentage of SARS-CoV-2 isolates carrying the alleles with D614G mutation in each period of time. **D.** Frequencies of D614G mutation in the S gene in each period of time. *p < 0.05; **p < 0.01.

### D614G mutation of S gene may destabilize S protein trimer and promote receptor binding and membrane fusion

The positive selected D614G mutation might play an important role for the adaptability of SARS-CoV-2 in both the host and the virus population[42]. Another explanation is that the mutation is driven by specific interaction between high level of virus sequence divergence and polymorphic host receptors or interacting proteins[43]. S protein is the key determinant for the tissue tropism and host range and specificity of coronavirus such as SARS-CoV-2. The virus infects host cells through the interaction between the S protein and its cellular receptor, named ACE2 [8]. In this process, virus entry requires the precursor S protein cleaved by cellular proteases including trypsin, furin, transmembrane serine protease 2 (TMPRSS2), or endosomal cathepsin L, which generate the receptor binding subunit S1 and the membrane fusion S2 [44-46]. From structural studies in both SARS-CoV and SARS-CoV-2, receptor binding domain (RBD) located at the C-terminal of S1 and the adjacent N-terminal domain (NTD) are relatively flexible, which is the feature required for receptor recognition and subsequent membrane fusion[47, 48]. We found that the D614G mutation is located at the subdomain 2 (SD2) that at the C-terminal of RBD and close to the two potential cleavage sites between S1 and S2 [48] (Fig 7A). Considering that positive selection is usually beneficial to the survival of the individual carrying the mutation, we speculate that the D614G mutation may facilitate structural conformation change to promote receptor binding or membrane fusion[5, 44], and in turn improving the infection efficiency. From the latest cryo-electron microscopy (cryo-EM) structure of SARS-CoV-2 S protein, the negatively charged sidechain of D614 points towards the positively charged sidechain of K854 from the neighboring monomer (Fig 7B) [48]. The distance between the closest atoms of the two residues is 2.6 Å, which is an optimal distance to form salt bridge (Fig 7C). From the modelled structure with D614G mutation, the distance is increased to 5.2 Å (Fig 7D), which would potentially abolish the salt bridge and destabilize the integrity of the S trimer in wild type. It has been reported that human receptor ACE2 binds to an “open” conformation of S protein, where RBD move away from the core structure and expose its receptor binding surface. The entire S trimer then undergoes a serial of dramatic conformation changes, including cleavages between S1 and S2, disassociation of S1 and post-fusion transformation of S2 [49, 50]. Changes including mutations at cleavage sites and adding internal crosslinks in S trimer would keep the protein in a stable and “closed” conformation where the receptor binding surface of RBD is inaccessible [48, 51]. Therefore, we hypothesize that the highly transmissible D614G mutation driven by the positive selection through evolution promotes accessibility of RBD by losing a critical salt bridge between the S protein monomers, which subsequently triggers membrane fusion upon ACE2 binding.

**Figure 7.**
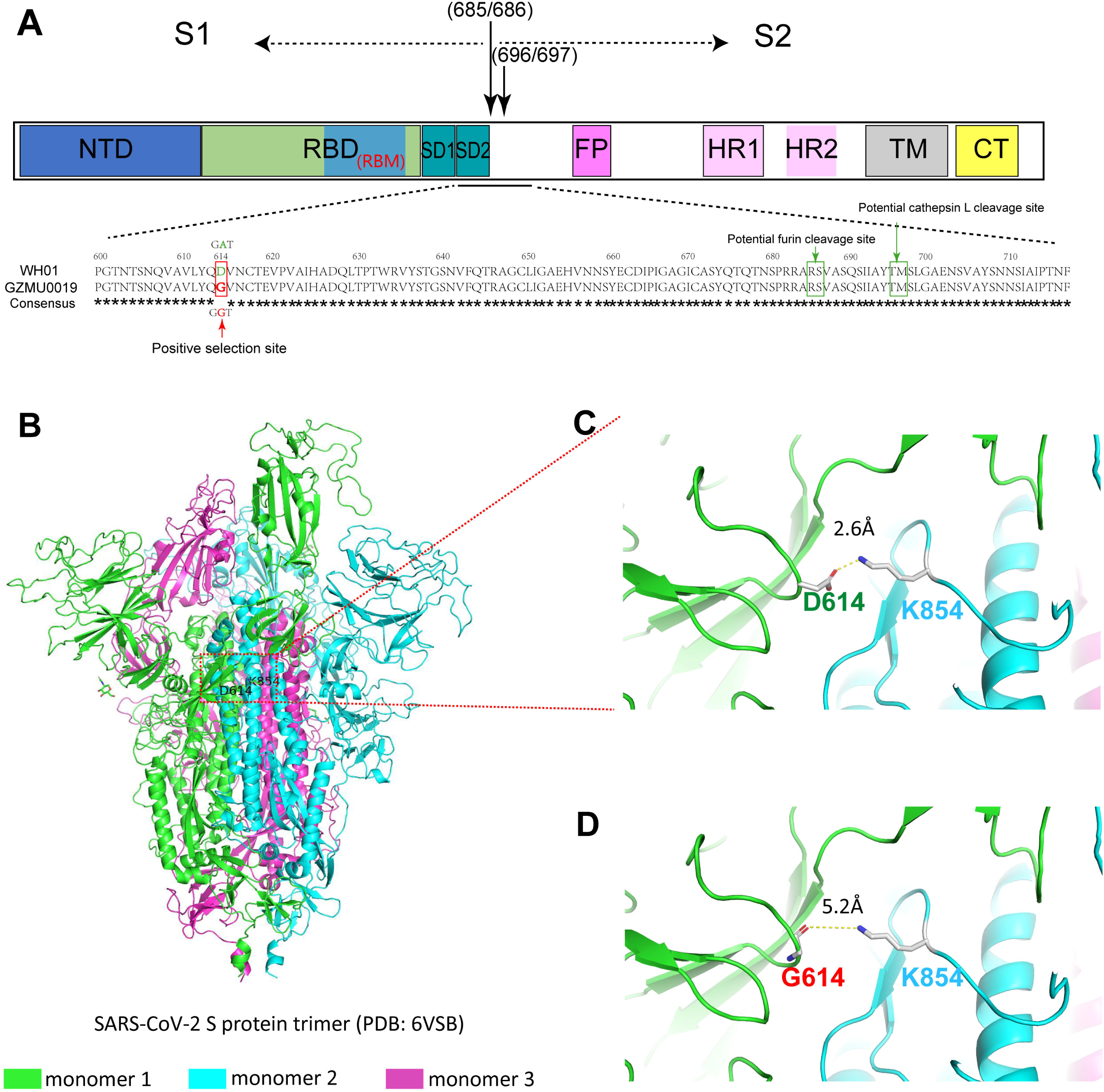
The structure of the S protein of SARS-CoV-2 and potential influence of D614G mutation on its structural change. **A.** Schematic of the primary structure of SARS-CoV-2 S protein colored by domains. Some boundary-residues are listed. The S1/S2 cleavage sites are indicated by arrows. RBD: receptor binding domain; RBM: receptor of binding motif; FP: fusion peptide, HR1/2: heptad repeat 1/2; TM: transmembrane domain; CT: cytoplasmic tail; NTD: N-terminal domain; CTD: C-terminal domain; SD1: subdomain 1; SD2: subdomain 2. The structure of the S protein trimer of SARS-CoV-2 and potential influence of D614G mutation on its structural change. **B.** Experimentally determined structure of SARS-CoV-2 S protein trimer (PDB ID is 6VSB and the amino acid sequences is the same as WH01 isolate). **C.** D614-K854 inter-monomer salt bridge. **D.** G614-K854 inter-monomer salt bridge. The distance of the salt bridge is increased from 2.6 to 5.2 Å in D614G mutation as shown.

## Conclusions

We present modern molecular evolution analyses on a large and comparative set of SARS-CoV-2 structural gene sequences, derived from an international collection of SARS-CoV-2 isolates. Distinct phylogenetic patterns of four structural proteins of SARS-CoV-2 are depicted. Protein sequence comparisons show E and M genes exhibit a relatively close relationship to bat SARSr-CoV, suggesting the evolution conservation of these two genes. In contrast, relatively high genetic variation is observed in N and S proteins among SARS-CoV-2 isolates, implying extensive adaptability of N and S genes. No clear intragenic recombination is detected of these four genes, suggesting that it is not the major force to drive the evolution of the four genes. However, our analyses show purifying selection pressure may be the main force operating the evolution at whole gene levels of SARS-CoV-2 during its human to human transmission. We also identify a codon in S gene definitively experiencing positive selection pressure, and always leads to the D614G mutation in S proteins. S alleles with D614G mutation have expanded rapidly among SARS-CoV-2 isolates. D614G mutation significantly extends the distance between monomers in the S protein trimer, which may disrupt the salt bridge formed by D614 and K854 between monomers, promote RBD opening, and facilitate the entry of the virus into host cells, thus contributing to the diffusion of this mutated alleles. Codon 5 of S gene is another potential positive selection site. Although a limited number of alleles with L5F mutation is identified, it may potentially affect the assembly and secretion of SARS-CoV-2. A close eye on L5F mutation may be required in case another expansion occurs. As S protein is a key target for SARS-CoV-2 vaccines, therapeutic antibodies, and diagnostics, the D614G mutation of S should be paid more attention. Owning that the exact mechanism remains unclear, further study should focus on the exact function of these mutation sites and how they affect the expansion of these mutated alleles on SARS-CoV-2.

## Acknowledgements

This research was supported by National Natural Science Foundation of China (grant number 31870001) to X.Y.Z.

## Supporting information

**S1 Table.** SARS-CoV-2 isolates information.

**S2 Table.** Log-likelihood values and parameter estimates for the SARS-CoV-2 N gene sequences. **S3 Table.** Detailed information of SARS-CoV-2 isolates with full length sequence of S gene. The data are organized by weekly.

**S4 Table**. Detailed information of SARS-CoV-2 isolates with full length sequence of S gene. The data are organized by panels. Each panel contains 200-300 isolates by combining isolates from several days.

**S1 Fig. The evolutionary relationship of N alleles. A.** Phylogenetic tree of N gene based on nucleotide sequences of 131 alleles. The evolutionary history is inferred using the Maximum Likelihood method and Tamura-Nei model. The tree is drawn to scale, with branch lengths measured in the number of substitutions per site. Bootstrap values more than 0.5 are shown. **B.** Parsimony network of SARS-CoV-2 N gene haplotype (allele) diversity obtained from 3090 isolates worldwide. Each oblique line linking between haplotypes (haplotype name is shown as its representative isolate name) represents one mutational difference. The ancestral haplotype, or root of the network, is labeled with a square, and represent haplotype name is marked red.

**S2 Fig. Evolutionary relationship of S alleles with or without L5F mutation. A.** Phylogenetic tree of S gene based on nucleotide sequences of 173 alleles. Each clade is highlighted with different color. Alleles are shown with their representative isolate names, and alleles with L5F mutation are highlighted in blue. Bootstrap values more than 0.5 are shown. **B.** Parsimony network of SARS-CoV-2 S gene haplotype (allele) diversity obtained from 3090 isolates worldwide. Each oblique line linking between haplotypes (haplotype name is shown as its representative isolate name) represents one mutational difference. Unlabeled nodes (Gray circle) indicate inferred steps have not found in the sampled populations yet. The ancestral haplotype, or root of the network, is labeled with a square, and represent haplotype name is marked green or red. The blue nodes indicate haplotypes with L5F mutation. Dotted boxes indicate major haplotype groups. Haplotypes include in red dotted boxes are with D614G mutation while those included in black dotted boxes are without D614G mutation.

**S3 Fig. Positive selection analysis of S gene codon 5 by Evolutionary Fingerprinting method.** Log (Bayes Factor) for positive selection at codon 5 of S gene and its frequencies. The cut-off value for the Bayes factor (BF) in the Evolutionary Fingerprinting method was set at 25 to reflect a positive selection at a given site (Posterior probability>0.95). Pr {BF>25} indicates posterior probability of Bayes Factor >25.

